# Modelling the effects of leaky predator-exclusion fences and their surrounding halo

**DOI:** 10.1101/737924

**Authors:** Kanupriya Agarwal, Michael Bode

## Abstract

Terrestrial fauna of the southern hemisphere, particularly Australia and New Zealand, have suffered significant declines and extinctions due to predation by introduced red foxes *Vulpes vulpes* and cats *Felis catus*. Predator-exclusion fences offer protection to these threatened species and allow their populations to persist and even flourish within their boundaries. These fences have traditionally been designed to stop the movement of both the invasive predators (into the fence), and the native animals (out of the fence). However, recent theory and evidence suggest that when native animals are able to move across the fence, they can create a population beyond the fence boundary. This phenomena has been called a “halo effect”, and has the potential to both expand the direct and indirect benefits of predator-exclusion fences, and to reduce their negative effects. However, the conditions under which such an effect can be achieved are uncertain. They include questions about which native species could support a meaningful halo, what levels of predation outside the fence can be tolerated, and how permeable the fence would need to be. Here, we formulate this problem as both a simple two-patch model and a spatial partial differential equation model. We use the two approaches to explore the conditions under which a halo can deliver conservation benefits, and offer clear insights into the problem.

## INTRODUCTION

Since the arrival of colonists from Europe, the terrestrial fauna of the southern hemisphere has experienced enormous population reduction and extinction (Boyer 2008; Miskelly et al. 2008; Radford et al. 2018; Woinarski et al. 2015). The fauna of ecosystems without native mammalian predators (principally Australia and oceanic islands) have been particularly decimated. European red foxes *Vulpes vulpes* and feral cats *Felis catus* bear a large portion of the responsibility (Legge et al. 2018; Nogales et al. 2004). In addition to the burden of predation they impose on native species, foxes and cats also amplify the negative impacts of habitat degradation, changing fire regimes, invasive herbivores, new diseases, climate change, and each other (Dickman 1996; Glen and Dickman 2005; Smith and Quin 1996; Woinarski et al. 2015). In Australia for example, thirty ground-dwelling mammal species have already been driven to extinction, but this is an ongoing catastrophe; at least 68 surviving species are highly susceptible to foxes and cats, and are at potential risk of extinction (Radford et al. 2018).

Conservation managers have devised a wide and inventive set of tools to reduce the effects of foxes and cats on threatened native species, including poison baiting, shooting, trapping, fumigation, and guardian animals. One of the most successful has been the construction of predator-exclusion fences; tall, wired, electrified fences, within which all invasive predators are removed. Australia increasingly relies on fenced enclosures to protect its native species. Alongside predator-free islands, these are the only action that can protect the 33 Australian mammal species that are classified as extremely susceptible to predation (Radford et al. 2018). Currently, there are 17 functional predator-exclusion fences across Australia protecting 25 distinct threatened mammal species including the bilby *Macrotis lagotis* and Gilbert’s potoroo *Potorous gilbertii* (Legge et al. 2018). Fences are also common in New Zealand, and are used across some islands of the Pacific (Somers & Hayward 2011).

Predator-exclusion fences do have limitations. They are expensive to construct and maintain (Bode et al. 2012), they can experience expensive and sometimes catastrophic failures (Bode and Wintle 2010; Helmstedt et al. 2014), and the predator-free environment they create may lead to behavioural and evolutionary maladaptation (although empirical results are mixed; Bannister et al. 2019; Hayward and Kerley 2009). However, their primary limitation is spatial: the largest predator-exclusion fence in Australia encloses 123 km^2^, a tiny proportion of the predator-filled surrounding landscape (Legge et al. 2018). Moreover, this limitation creates one of their largest challenges. Native animal populations are sometimes so successful inside fences that they reach unsustainable levels (Moseby et al. 2018). Australian predator exclusion fences, most of which date from the 2000s and are sited in fragile semi-arid and arid environments (Legge et al. 2018), are increasingly reporting high densities of particular herbivores. These elevated populations are damaging the local vegetation, with negative consequences for other threatened species that have similar diets or habitat requirements (Linley et al. 2017; Moseby et al. 2018).

One simple solution to overpopulation within an exclusion fence is to release the surplus animals into the local landscape, outside the fence. Predator-exclusion fences are designed to be “perfect”, with neither predators or prey crossing their boundary. Instead, they could be allowed to be “leaky”, where threatened species can emigrate outside the fence. These individuals would spread from the vicinity of the fence, extending the native population and its benefits into a region that could be much larger than the fenced area. While these released animals would experience high predation rates, the constant emigration of surplus individuals from the fenced population could potentially maintain the outside population. The flow of individuals across the fence would be balanced by the population decline from predation, creating an anthropogenic stable, source-sink system (Pulliam 1988). The source portion of the system would be the “halo” population around the fence, extending out from the high densities inside the fenced area, eventually declining to zero (Figure 1).

**Figure 1:**
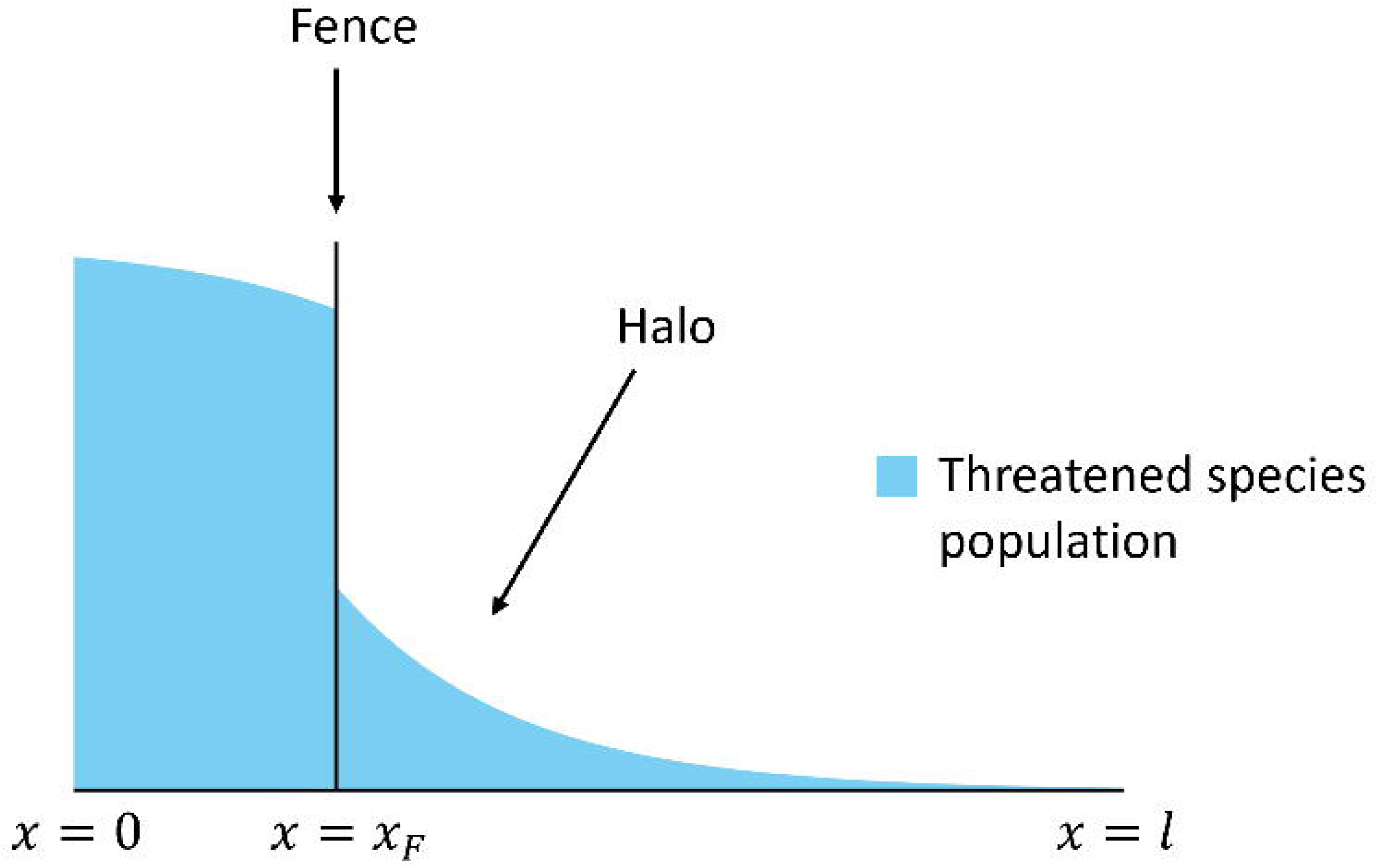
Schematic diagram of a partially fenced landscape, with a surrounding halo caused by a leaky fence at x = x_F_. The height of the blue shaded region denotes the density of a native animal, which is greater inside the fence (x < x_F_), and declines to zero at large distances (x ≫x_F_), where predation causes a net decline of the species. The bounded domain of the model extends to l ≫x_F_. Note that because the model structure is symmetric about x = 0 (i.e. there is another fence at x − x_F_) we only show the right-hand part of the domain.

**Figure 1a:**
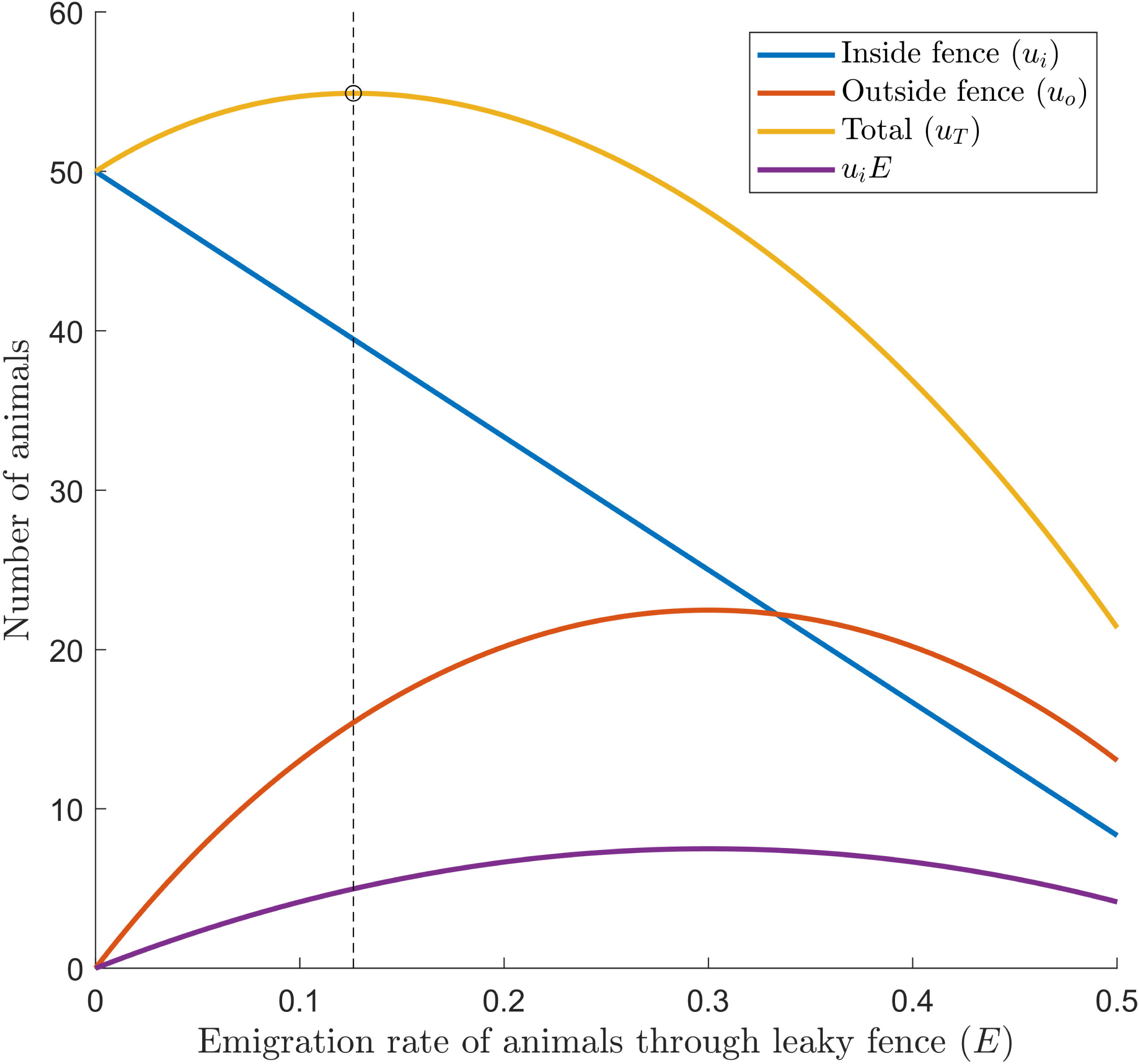
Figure 1: Equilibrium solutions of Eq. (1) for a range of values of E (shown on the x-axis). Other parameter values were chosen to be r = 0.6, α = 0.9, P = 1, k_i_ = 50 and k_0_= 400. Lines indicate the equilibrium populations inside the fence (blue), outside the fence (orange), and overall (yellow). The number of individuals emigrating per timestep is shown in purple. The vertical dashed line shows the optimal solution that maximises the total abundance.

Halos around predator-exclusion fences have already been observed for populations of falcon *Falco novaeseelandiae* and New Zealand pigeon *Hemiphaga novaeseelandiae* in New Zealand (Russell et al. 2015), although these halos were created automatically: the birds’ ability to fly allows them to move across the boundary at will to nest or forage in the wider landscape. Conservationists in Wellington are hoping that eco-sanctuaries and invasive predator management can deliver a similar halo effect for the threatened kiwi species *Apteryx* spp. (Burns et al. 2012; Roy 2018). In Australia, fences are already trialling the use of species-specific one-way gates to regulate populations within multi-partition fences, with plans to allow the one-way movement of over-abundant herbivores into the surrounding landscape (Butler et al. 2019; Crisp and Moseby 2010). Although not associated with a fence, halo effects have also been observed in Western Australia rock wallaby populations *Petrogale* spp. that are protected from feral predation by poison baiting (Glen et al. 2013; Kinnear et al. 2015). Baiting increased the rock wallaby population within the treated area, and by increasing the frequency of baiting, the threatened animals were able to thrive both in the baited areas and its surrounding, unbaited areas (Kinnear et al. 2015).

In this paper, we use theoretical spatial population models to explore and offer insights into the dynamics of leaky predator-exclusion fences and halos. Our goal is to understand the conditions under which a leaky fence will deliver a net benefit to a threatened species, to estimate how many animals should be released from (or should be allowed to naturally leave) the fenced region, and to estimate how large the halo effect will be – both in terms of its spatial extent and the size of the population inside and outside the fence.

## TWO-PATCH MODEL

At its most basic level, a leaky predator-exclusion fence can be described by a two-patch source-sink model, with the first patch representing the source population within the predator-exclusion fence, the second patch representing the sink population outside the fence, and with one-way dispersal between the two patches representing the movement of the threatened species through the fence. We describe this simple system with a coupled pair of harvested logistic models (Murray 1989):

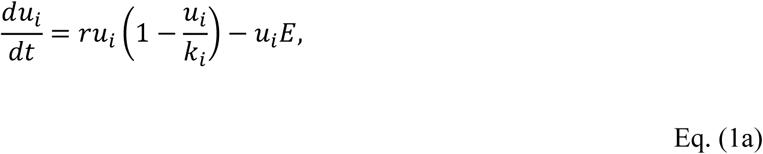

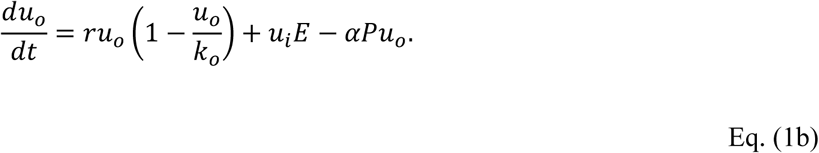

where *i* and *o* denote the patches inside and outside the fence respectively, *u*_*x*_ is the native species’ abundance in each patch, *r*_*x*_ is its per-capita growth rate, and *α* is the predation rate. The parameters *k*_*i*_ and *k*_*o*_ denote the maximum possible abundance of the native species inside and outside the fence. The maximum abundance inside the fence has a straightforward interpretation – the carrying capacity density of the landscape multiplied by the total fenced area. The total area outside the fence is more complicated, since we don’t yet know the spatial area occupied by the halo. For the two-patch model we will simply assume values where *k*_*o*_ ≫*k*_*i*_, and return to this question more explicitly in the spatial model. The parameter *P* indicates the number of predators in the outside patch, which we assume to be constant through time. The parameter *E* describes the rate of emigration across the fence – essentially, how leaky the fence is – measured by the per-timestep probability that an animal inside the fence will emigrate.

In the models that follow, we will not consider species that can persist without a fence i.e. when *r* > *αP* (see *Supplementary Methods* for more details), since we assume that fences will only be designed for the set of species that need them most. We also assume that management objective is to maximise the total abundance of the threatened species – both inside and outside the fence:

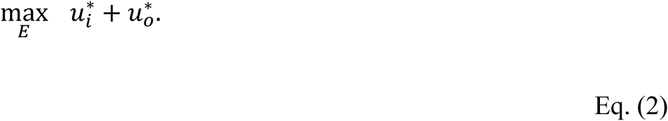

Calculating the solution to Eq. (1) is a two-step process. First, we solve for the equilibrium abundance on the inside of the fence; then, we use this interior solution to solve for the equilibrium abundance outside the fence. At equilibrium, the total population of the threatened species is then:

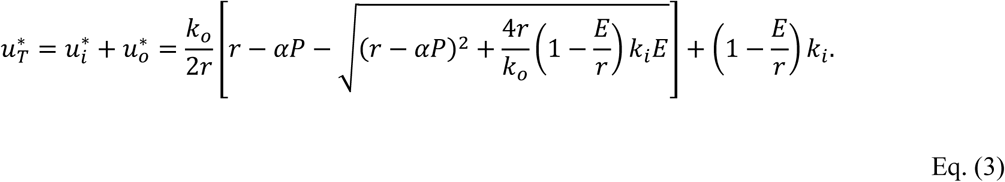

To find the optimal emigration rate in Eq. (2), we maximise 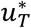 with respect to *E* (verifying that the second derivative is negative), giving:

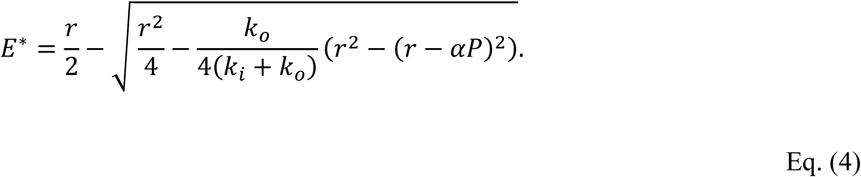

Figure 2 shows this result graphically. The form of this solution suggests that the optimal emigration rate depends on a contrast between the threatened species’ intrinsic growth rate (the term *r*^*2*^), and its growth rate in the presence of the predator outside the fence *(r- αP)*^*2*^. The optimal emigration rate should also increase as the relative area outside the fence *k*_*o*_*/(k*_*o*_ *+ k*_*i*_) increases. Lower density populations (spread across larger outside areas) have higher per-capita growth rates, encouraging higher optimal emigration rates.

**Figure 2:**
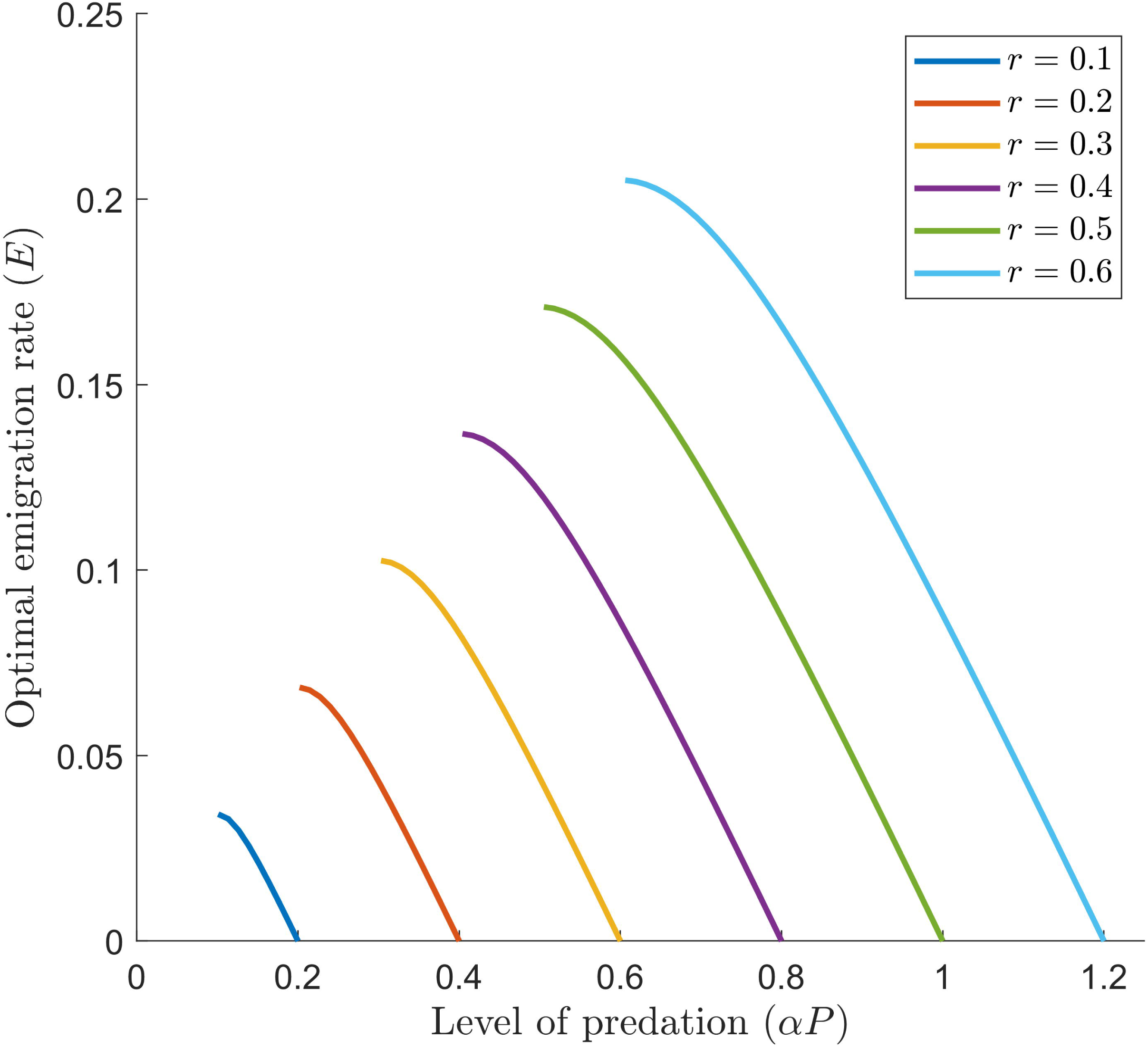
Optimal rate of emigration of native animals across the fence (i.e. how leaky the fence should be) for different levels of predation. Coloured lines represent different per capita growth rates of the threatened species.

The blue line in Figure 2 indicates that the number of animals inside the fence declines linearly as the emigration rate increases. When the fence is perfect (i.e., *E* = 0), the red line shows that the outside population is zero, since this species cannot persist in the presence of invasive predators at density *P*. As the emigration rate increases, the outside population at first increases rapidly, slows, then eventually declines. The decline occurs because the increasing values of *E* are delivering a larger proportion of a much smaller population (purple line). The total population (inside and outside) also increases and then decreases, with the maxima occurring before the outside population is maximised.

In Figure 4, we show the optimal emigration rate, given by Eq. (4), for different values of predation and growth rate. For *r =* 0.1, for example, the species doesn’t require a fence while *αP <* 0.1, since it can persist in the presence of predation. At higher levels of predation, the species’ total population will benefit from a leaky fence, but once the level of predation exceeds double the growth rate, a perfect fence is needed (once the curves intersect the *x*-axis). The coloured lines show that populations with higher growth rates can support higher emigration rates.

**Figure 4:**
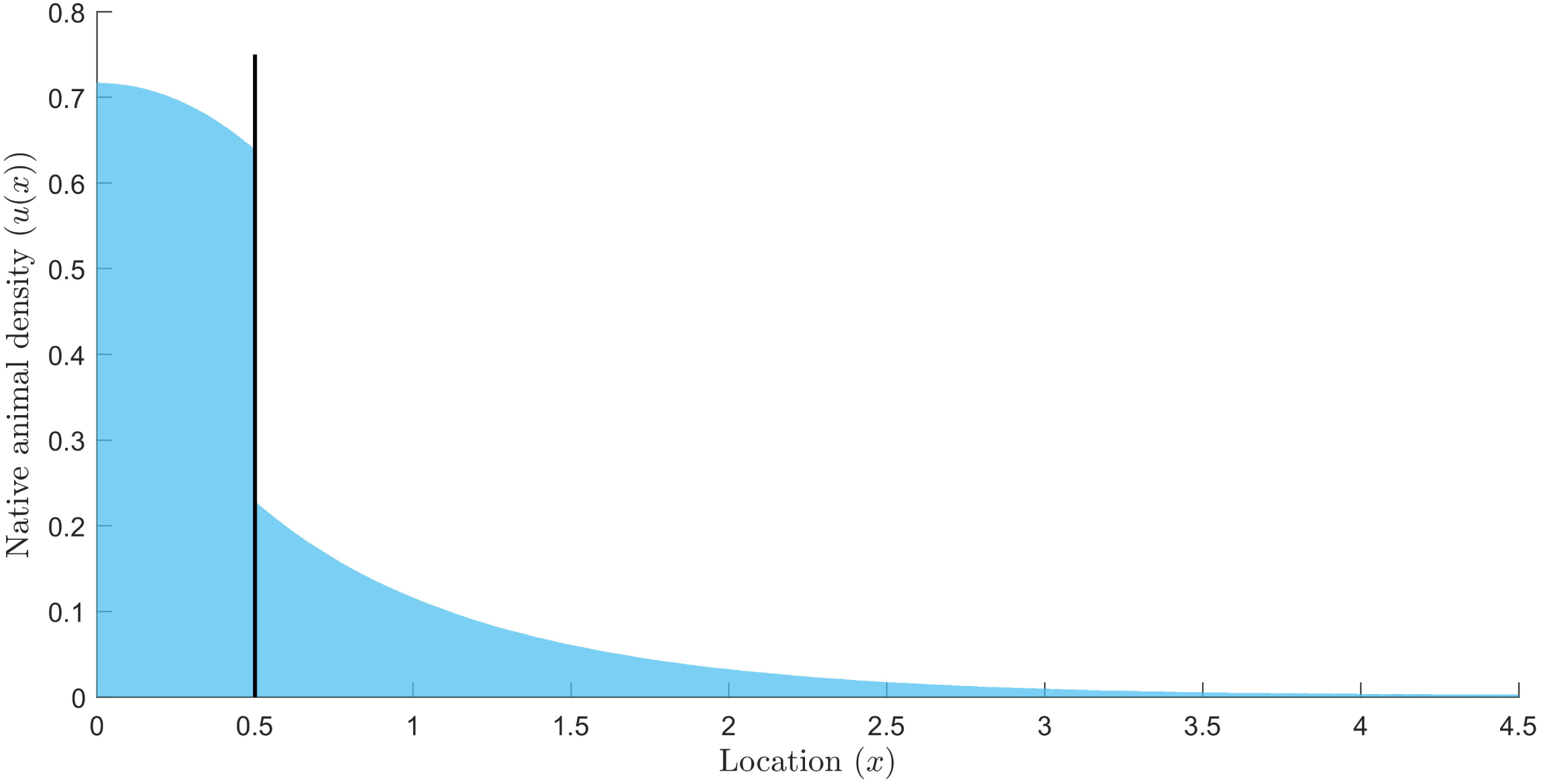
Example solution to Eq. (5); the equilibrium density distribution of the threatened species population, given the parameter values r = 0.6, α = 0.9, P = 1, k = 1, D = 0.3 and E = 0.15.

## FISHER-KOLMOGOROV MODEL

The two-patch model can offer insights into the optimal emigration rate, but such a simple representation of space cannot be used to describe the size or shape of the halo, nor consider how the movement of the threatened species through the landscape could alter the halo. We therefore reformulate the problem using a one-dimensional partial differential equation, modified from the well-known Fisher-Kolmogorov (FK) equation (Murray 1989, Okubo 1980):

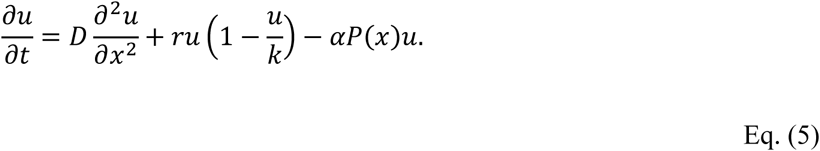

We assume that the landscape shown in Figure 1 is homogeneous, with constant diffusivity *D*, and maximum threatened species density *k*. The first term on the right-hand side of Eq. (5) describes the random movement of animals in the landscape due to diffusion, the second term describes the logistic growth of the prey species, and the third term describes predation. We must also define the predator density *P(x)* as a piece-wise function that is 0 inside and at the fence:

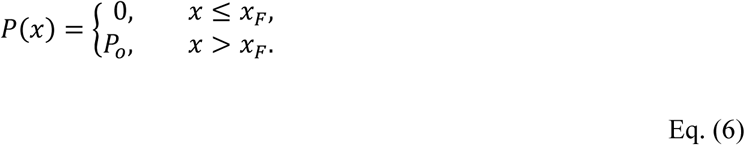

The threatened species’ abundance outside the fence is influenced by its abundance inside the fence, but not vice versa. Ecologically, this reflects the one-way exchange between the source fenced population, and the sink population outside the fence; mathematically, this allows us to solve the outside equation by substituting the interior equilibrium abundance into the outside equation as a matching boundary condition at the fence location.

To determine the equilibrium density distribution, we first solve Eq. (5) for the fenced region (0≤ x≤ *x*_*F*_). In the interior of this region (at *x =* 0), the density profile is symmetric, and we therefore model the left-hand interior boundary condition as a Neumann reflecting boundary condition. The right-hand boundary condition (at *x = x*_*F*_) is also a Neumann condition, where the rate of movement across the leaky fence is equal to the emigration rate *E* multiplied by the number of animals inside the fence. We then solve the system for the outside region (*x*_*F*_ ≤*x* ≤ *l*), where the left-hand Neumann boundary condition at *x*_*F*_ is defined by the equilibrium solution of the interior population. These boundary conditions ensure the conservation of animals (i.e., every individual that exits the fenced population enters the halo population). More details on the solution are given in the *Supplementary Methods*.

To form the density distribution profile, Eq. (5) was numerically solved using a finite difference method. An example solution is shown in Figure 4.

**Figure 5:**
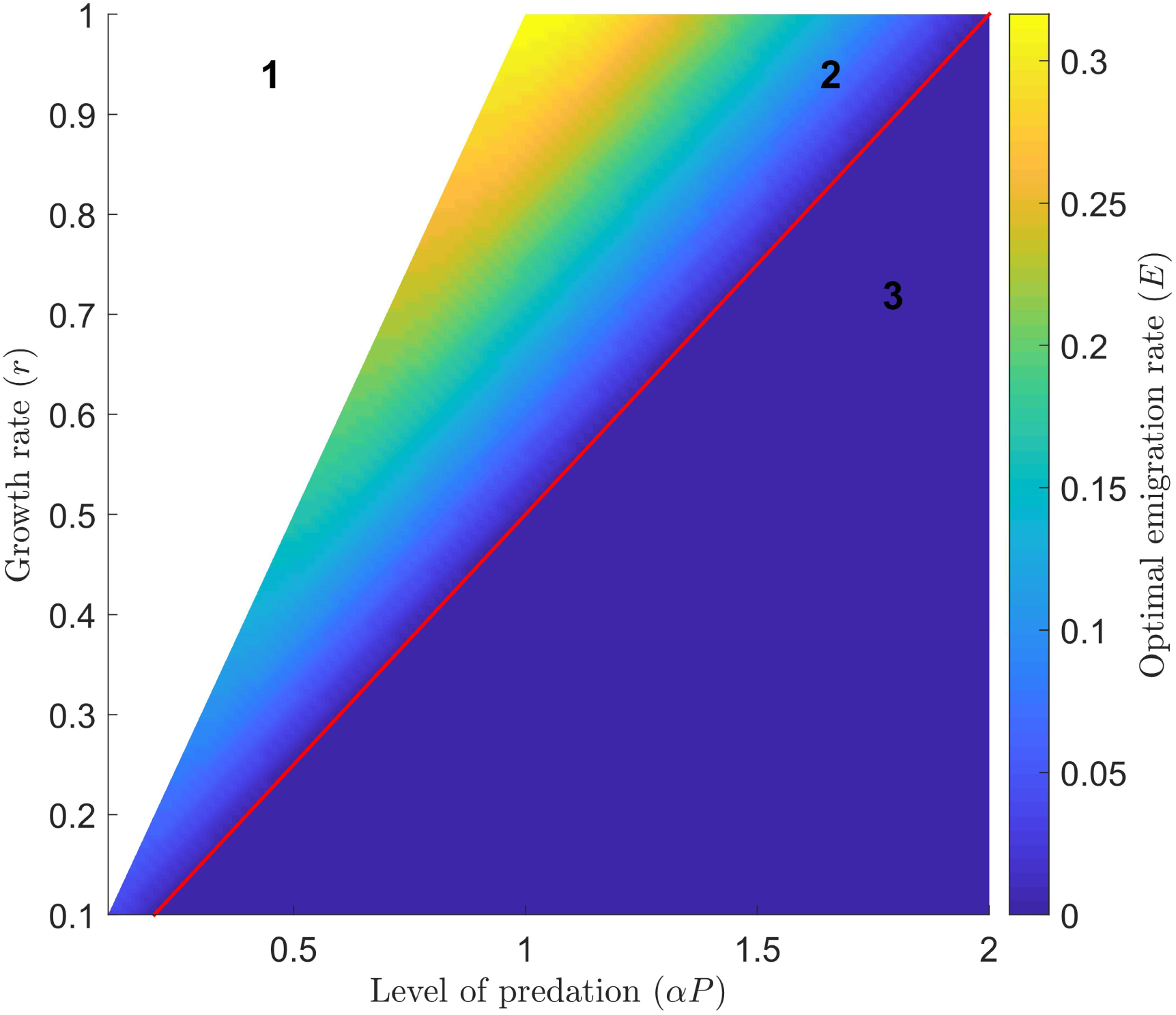
Optimal emigration rate (percentage of animals leaving the fence) for the Fisher-Kolmogorov model given the parameter values k = 100 and D = 0.5. There are three categorically different optimal solutions: no fence (section 1), imperfect fence and halo (section 2), and perfect fence (section 3).

Depending on the parameters chosen, three qualitatively different solutions maximise total abundance. The first category is species that can persist in the presence of predation – we assume that these species do not require a fence at all. The second category is species that need a perfect fence. For these species, any animals that move outside the fence are killed before they can effectively reproduce, so emigration only decreases the total population size. The third category of species benefit from leaky fences. These species cannot persist outside the fence, but they can still reproduce outside the fence to some extent. Their emigration therefore contributes positively to the overall population. For species that need fences (i.e., the second and third categories), the optimal rate of escape from the fence decreases as the level of predation increases. Once again, if the level of predation exceeds double the growth rate, the optimal decision is to construct a perfect fence.

Our solution of interest – when the leaky fence is able to produce a halo that increases the total population size – is shown in Figure 4. There is an abundance of animals inside the fence, with the density highest at the interior of the fence. The interior density profile curves, as a consequence of emigration across the leaking fence. This species cannot persist given the level of predation outside the fence, as seen in the exponentially declining tail on the right hand of Figure 4, but it can maintain a positive equilibria halo, far beyond the fence boundary. At the location of the fence, there is a discontinuity in the density function. Here, the gradients are equal: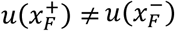, but the Neumann boundary conditions ensure that the first derivative is continuous: 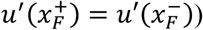.

For the same parameter values, the two-patch and Fisher-Kolmogorov formulations of the model are in general agreement. For the values shown in Figure 2 for example, the optimal rate of emigration of animals through the fence is *E*^*^ ≈ 0.13 for the two-patch model, and the slightly larger value of *E*^*^≈ 0.12 for the Fisher-Kolmogorov model (see *Supplementary Methods*). This means that the total population would be maximised if approximately 12-13% of animals from the fence were released into the surrounding area, at each timestep.

For a range of growth rate and predation rate parameters, we can calculate the optimal emigration rate for the Fisher-Kolmogorov formulation. The results, shown in Figure 5, reveals the three categories of solution. As explained earlier, species that can survive in the presence of predation do not need a fence (section 1). At the other end of the spectrum, a perfect fence (section 3) with a zero emigration rate is optimal for particularly predator-susceptible species. In the middle, a leaky fence (section 2) has a positive optimal emigration rate that allows some animals to leave, and creates a halo effect beyond its boundaries. As the red line suggests, the transition between a perfect fence and a leaky fence occurs when the level of predation exceeds double the growth rate.

## DISCUSSION

To determine whether leaky fences are a viable option for threatened species, we first formed a two-patch model, which treats the inside of the fence and fence’s surrounding areas as two separate regions. We found an analytical solution to the two-patch model, which offered some insight to the behaviours of the optimal solution. To incorporate the spatial aspect into the model, we then formed the FK model using a spatial PDE. We found that some threatened species need leaky fences and halos to maximise their population, whilst some threatened species need perfect fences as they cannot survive in the presence of predation at all. In general, we saw that for species that need fences, the optimal emigration rate from the fence decreases as the level of predation increases. For example, for a threatened species with given growth rate, if the level of predation in a certain area is greater than the growth rate, but less than two times the growth rate (*r* ≤ *αP* ≤ *2r*), then the species will benefit from a leaky fence. Otherwise, if the level of predation is greater than two times the growth rate, then the threatened species will no longer benefit from a leaky fence, and would be better off with a perfect fence.

Our results are all designed to maximise the total population of the threatened species, inside and outside the leaking fence. There are many other legitimate conservation objectives that do not depend on the total abundance, or that only partly depend on abundance. For example, managers might want to maximise the spatial extent of the threatened population – i.e., they might want a population both outside and inside the fence, to minimise the chance of a catastrophe extirpating the whole population. These results could be calculated directly from our model – although only the spatial FK model – using different formulations of the objective function Eq. (2). Under this objective, the range of conditions under which a leaky fence is optimal would be wider than under the objective we use in these analyses.

Our model does not address the abundance of predators outside the fence. But baiting would effectively reduce predator populations and improve the threatened species populations in the areas surrounding the fence. For species that we have identified as requiring perfect fences, baiting the surrounding regions of the fence and allowing them to leave the fence could provide them with extra protection from predators and thus allow their population to grow in the wild, beyond the fences and baited areas.

In pursuit of theoretical insights, important elements of realism were stripped from out two models. We treated the diffusivity of the landscape to be homogeneous, but this does not reflect the landscape for the inside of a fence or its surrounding area. We also defined the number of predators to be constant outside the fence, but predator numbers fluctuate as a consequence of environmental and demographic stochasticity, and will also respond to the prey density. While this means that the input of new prey individuals from the fenced population could potentially attract new predators, fox and cat abundances in Australia respond to invasive herbivore densities (primarily rabbits *Oryctolagus cuniculus*), which are generally much higher than those of native species (Southgate and Masters 1996). Finally, our parameter values were chosen to illustrate particular categories of solution, rather than particular real conservation contexts. We cannot therefore draw specific conclusions about where leaky fences will work, and for which species.

