## Supplementary Methods for "Modelling the effects of leaky predator-exclusion fences and their surrounding halo"

In the supplementary methods, we provide full derivations for three conclusions drawn in the main text, and an additional figure that compares the equilibrium solution of the FK model with the two-patch model.

### 1. Species that can and cannot persist without fences

A common model of a predator-prey system is,

$$\frac{du}{dt} = ru\left(1 - \frac{u}{k}\right) - \alpha P u$$

Eq. (S1)

where $u$ is the total number of prey, $r$ is the per capita growth rate of prey, $k$ is the carrying capacity of prey, $P$ is the number of predators, and $\alpha$ is the predation rate. The first term in Eq. (S1) is a logistical model which is a common description of population dynamics, and it represents the population growth of the prey. The second term represents the predator-prey interaction and the predation rate is essentially the number of predator-prey interactions that result in deaths.

To understand how the variables in this model (e.g. growth rate of prey, predation rate) affect the total number of prey, we can solve Eq. (S1). We want the solution at equilibrium, so we set Eq. (S1) to be 0 and then solve for $u^*$.

$$\frac{du}{dt} = 0 = ru\left(1 - \frac{u}{k}\right) - \alpha P u$$

Eq. (S2)

$$u^* = \frac{k(r - \alpha P)}{r}$$

Eq. (S3)

We shall now refer to the predation rate multiplied by the number of predators, or $\alpha P$ as the 'level of predation'. From the solution in Eq. (S3), we can see that if the growth rate is equal to the level of predation, the total number of prey will be 0, i.e. if $r = \alpha P$, then $u = 0$. Similarly, if $r < \alpha P$, then $u < 0$, however the number of prey cannot be negative, so we assume that it is actually 0. If the growth rate is greater than the level of predation, the total number of prey will be greater than zero. Thus, when $r > \alpha P$, the species can persist in the presence of predation.

So, when $r \leq \alpha P$, the prey species require predator-exclusion fences to conserve their population, but when $r > \alpha P$, the prey do not require fences and can co-exist with their predators. Now, this may not always be true when $r$ is only slightly larger than $\alpha P$, but we assume that it is true for our models and we present our solutions based on this.

### 2. Analytical solution to two-patch model

The total population of the endangered species is given by,

$$u_T^* = u_i^* + u_o^*.$$

Eq. (S4)

Since we are concerned with the solution at equilibrium, we set

$$\frac{du_i}{dt} = 0,$$

Eq. (S5a)

$$\frac{du_o}{dt} = 0.$$

Eq. (S5b)

Substituting (S5a) into Eq. (1a), we then solve for $u_i^*$,

$$0 = ru_i\left(1 - \frac{u_i}{k}\right) - u_iE,$$

Eq. (S6)

$$u_i^* = \left(1 - \frac{E}{r}\right)k_i.$$

Eq. (S7)

Then, substituting Eq. (S5b) and (S7) into Eq. (1b), and solving for $u_o^*$ we get,

$$0 = ru_o\left(1 - \frac{u_o}{k_o}\right) + \left(1 - \frac{E}{r}\right)k_iE - \alpha Pu_o,$$

Eq. (S8)

$$u_o^* = \frac{k_o}{2r}\left[r - \alpha P - \sqrt{(r - \alpha P)^2 + \frac{4r}{k_o}\left(1 - \frac{E}{r}\right)k_iE}\right].$$

Eq. (S9)

Thus, substituting Eq. (S7) and (S9) into Eq. (S4), we see the total number of animals is given by Eq. (3). To find the optimal emigration rate of animals through the fence, we must calculate which value of $E$ gives the maximum total number of animals. So, we find the derivative of $u_T^*$ with respect to $E$, and set it to 0 to solve for the optimal emigration rate.

$$\frac{du_T^*}{dE} = 0 = \frac{2E - r}{\sqrt{(r - \alpha P)^2 + \frac{4r}{k_o}\left(1 - \frac{E}{r}\right)k_iE}} - 1,$$

Eq. (S10)

The optimal rate of emigration is then given by Eq. (4).

#### 3. Boundary conditions of Fisher-Kolmogorov model

Eq. (5) was solved at equilibrium, and so we set

$$\frac{\partial u}{\partial t} = 0.$$

Eq. (S11)

Eq. (5) now becomes an ordinary differential equation,

$$0 = D\frac{d^2u}{dx^2} + ru\left(1 - \frac{u}{k}\right) - \alpha P(x)u.$$

Eq. (S12)

To solve this ODE, it can be rewritten as a system of first-order ODEs by setting $u = u_1$ and $u' = u_1' = u_2$:

$$\frac{du_1}{dx} = u_2$$

Eq. (S13a)

$$\frac{du_2}{dx} = \frac{1}{D}\left[P(x)u_1 - ru_1\left(1 - \frac{u_1}{k}\right)\right]$$

Eq. (S13b)

The boundary conditions for the fenced region, i.e. when $0 \leq x \leq x_F$ are:

$$\left.\frac{du_1}{dx}\right|_{x=0} = 0,$$

Eq. (S14a)

$$\left.\frac{du_1}{dx}\right|_{x=x_F} = -u_1 E.$$

Eq. (S15b)

The boundary conditions for the outside region, i.e. when $0 \leq x \leq x_F$ are:

$$\left.\frac{du_1}{dx}\right|_{x=x_F} = -u_1 E,$$

Eq. (S16a)

$$\left.\frac{du_1}{dx}\right|_{x=l} = 0.$$

Eq. (S16b)

When solving for the outside region, the solution to the inside region is used as the value of $u_1$ in Eq. (S16a). There is a discontinuity at the fence since we are matching Dirichlet conditions, Eq. (S17), instead of Neumann conditions, Eq. (S18).

$$\left.\frac{du_i}{dx}\right|_{x=x_F} = \left.\frac{du_o}{dx}\right|_{x=x_F},$$

Eq. (S17)

$$u_i(x_F) = u_o(x_F).$$

Eq. (S18)

In other words, the value at the fence is the rate of change of the solution inside rather than the value of the solution inside. Since we are matching Dirichlet boundary conditions, Eq. (S15b) is the same as Eq. (S16a).

### 4. Equilibrium solution for FK model with different emigration rates

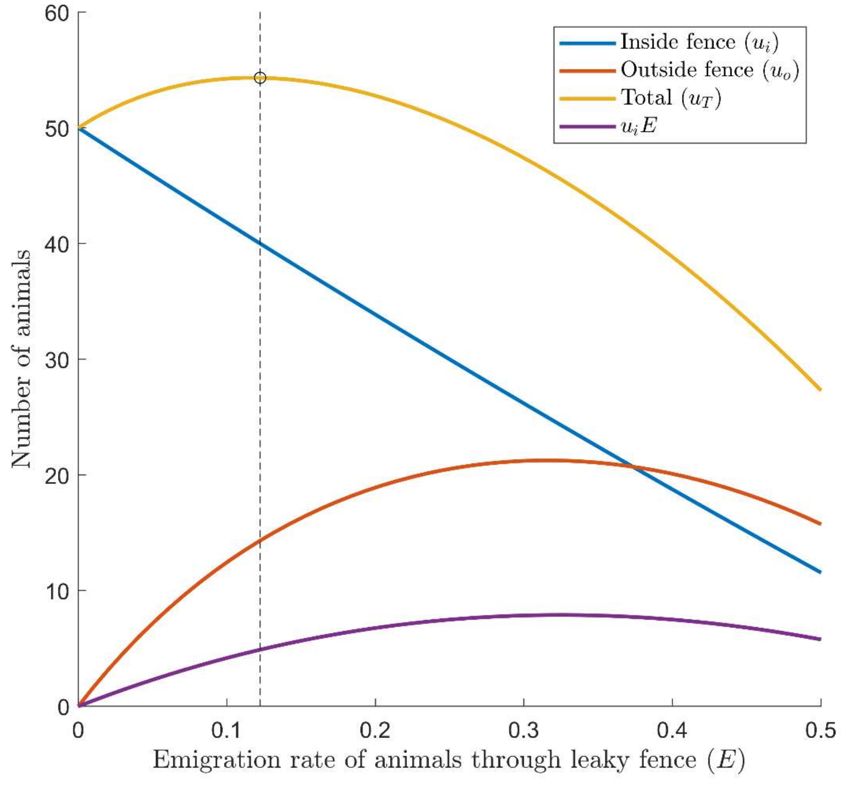


**Figure S1:** *Equilibrium solutions of Eq. (5) for a range of values of E (shown on the x-axis). Other parameter values were chosen to be* $r = 0.6$, $\alpha = 0.9$, $P_o = 1$, $k = 100$ *and* $D = 0.5$. *The coloured lines indicate the populations inside and outside the fence, overall, and the purple line denotes the rate of emigration. The vertical dashed line shows where the total abundance is maximised.*
